# The biogenesis of extracellular vesicles from *Staphylococcus aureus* and their application as a novel vaccine platform

**DOI:** 10.1101/255273

**Authors:** Xiaogang Wang, Christopher Weidenmaier, Jean C. Lee

## Abstract

Gram-positive bacteria secrete extracellular vesicles (EVs) that package diverse bacterial antigens and play key roles in bacterial pathogenesis. However, the mechanisms underlying EV production in Gram-positive bacteria are poorly understood. We purified and characterized EVs from a community-associated methicillin-resistant *Staphylococcus aureus* isolate (USA300) and investigated mechanisms underlying EV production. Native EVs contained 165 proteins, including cytosolic, surface, and secreted proteins, autolysins, and numerous cytolysins. Staphylococcal alpha-type phenol-soluble modulins (surfactant-like peptides) promoted EV biogenesis, presumably by acting at the cytoplasmic membrane, whereas peptidoglycan crosslinking and autolysin activity were found to increase EV production by altering the permeability of the staphylococcal cell wall. To address the immunogenicity of EVs, we created engineered EVs (eng-EVs) by expressing detoxified proteins Hla_H35L_ and LukE in EVs generated from a nontoxic *S. aureus ΔagrΔspa* mutant. Eng-EVs exhibited no cytotoxicity in vitro, and mice immunized with the eng-EVs produced toxin-neutralizing antibodies and showed reduced lethality in a mouse sepsis model. Our study reveals novel mechanisms underlying *S. aureus* EV production and highlights the usefulness of EVs as a novel *S. aureus* vaccine platform.

## Introduction

*Staphylococcus aureus* is a pathogenic bacterium that causes a wide range of human infections, ranging from mild skin lesions to more invasive and life-threatening infections, such as bacteremia, endocarditis, wound infections, and pneumonia. The pathogenesis of *S. aureus* infections is generally attributed to a wide array of virulence determinants including surface proteins^1^ and glycopolymers^2^, as well as multiple secreted proteins, such as cytolysins, superantigens, leukotoxins, hemolysins, and proteases^3^. Although several specific export pathways have been described in *S. aureus*, the secretome often includes proteins that lack export signals and have typical cytoplasmic functions. The mechanisms by which cytoplasmic proteins are excreted by *S. aureus* has attracted recent interest^4^, ^5^, and there is increasing evidence that these proteins may be secreted within membrane vesicles^6-9^.

Secretion of extracellular vesicles (EVs) is a process common to eukaryotes, archae, and bacteria^10^. EVs are nano-sized, spherical, bilayered membrane vesicles with a cargo that includes diverse proteins, polysaccharides, nucleic acids, and lipids. EV formation by Gram-negative bacteria was first observed by electron microscopy in the 1960s^11,12^,and these bacteria secrete what are now referred to as outer membrane vesicles (OMVs). The generation of OMVs appears to occur by phospholipid accumulation in the outer leaflet of the outer membrane, followed by the formation of outer membrane protrusions that pinch off to form vesicles^13^. OMVs likely play important roles in bacterial pathogenesis due to packaging of multiple virulence factors^14^, and the ability of OMVs to serve as immune modulators by inducing innate and adaptive immune responses^15^.

More recent work has described the production and release of extracellular vesicles (EVs) from Gram positive bacteria, such as *S. aureu*^16,17^, *Streptococcus pneumoniae*^7^, and *Bacillus anthracis*^6^. Only actively metabolizing bacteria generate EVs, and EVs are not released by killed cells^9^. Due to the presence of a thick peptidoglycan (PGN) structure surrounding the bacterial cell, the extracellular release of EVs from Gram positive microbes is a complex process that is poorly understood. That EVs from Gram positive organisms also play important roles in host-parasite intereractions is supported by reports that EVs may contain biologically active toxins, exhibit cytotoxicity, and elicit pro-inflammatory mediators^9^. Additional findings indicate that EVs positive for *S. aureus* toxins elicit skin barrier disruption in mice with characteristic atopic dermatitis-like skin inflammation^18-20^, highlighting their potential contribution to *S. aureus* disease. However, the toxicity of staphylococcal EVs has until now hampered a relevant study of their immunogenicity and potential use as a novel vaccine platform.

Development of bacteria-derived vesicles as a multivalent vaccine platform is feasible since vesicles package an array of different antigens including those that are cytosolic, membrane-associated, secreted, and surface exposed. A vaccine containing *Neisseria meningitidis* recombinant proteins combined with group B OMVs was licensed to protect humans against meningococcal B disease in the U.S. and other countries^21^, attesting to the efficacy of this vaccine platform. Additionally, a growing number of studies involving the use of OMVs as vaccines against bacterial pathogens have shown protection in experimental infection models^22-25^.

Despite repeated efforts to develop experimental vaccines and immunotherapeutics against *S. aureus*, neither have proven effective in preventing staphylococcal infections in humans^26^. Mice immunized with native EVs from *S. aureus* ATCC 14458 responded with a robust T cell response and were protected against staphylococcal lung infections, although the cytolytic activity of EVs prepared from wild type (WT) *S. aureus* was not addressed in this study^27^. Similarly, EVs isolated from *S. pneumoniae* protected mice against lethal pneumonia^7^. Despite the documented immunogenicity and protective efficacy of bacterial EV-based vaccines, EV preparations derived from some bacterial pathogens may contain toxins or other virulence factors that potentially damage host cells^17,28-31^. The development of EVs as a vaccine platform will require a more thorough characterization of the mechanisms of EV biogenesis to allow for consistent production with adequate quality assurance.

In the present study, we generated, purified, and characterized EVs isolated from *S. aureus* USA300, a predominant CA-MRSA clone in United States, and investigated the biogenesis of EV formation. Our study reveals distinct mechanisms that facilitate EV production at multiple stages. Phenol soluble modulins (PSMs) act at the membrane level to facilitate vesicle budding at the cytoplasmic membrane, whereas cell wall porosity is modulated by PGN cross-linking and production of autolysins. We investigated the cytotoxicity and immunogenicity of staphylococcal EVs and explored their usefulness as a novel vaccine platform. By genetically engineering a nontoxic mutant *S. aureus* strain to over-produce detoxified alpha hemolysin (Hla_H35L_) and a leukocidin monomer (LukE), we created engineered EVs (eng-EVs) that were immunogenic, nontoxic, and protected against *S. aureus* lethal sepsis in mice. Our investigations will not only further the development of this novel vaccine platform, but also promote further studies of the impact of EVs on the pathogenicity of *S. aureus* and other Gram-positive pathogens.

## Results

### Isolation of EVs from *S. aureus* USA300 strain JE2

EV formation by at least 10 different *S. aureus* strains has been reported^16,17,28,32^, and EV shedding is likely common to many clinical isolates. We isolated and characterized EVs from USA300 strain JE2, representing the major clone associated with CA-MRSA disease in the U.S^33^. JE2 culture supernatants were filtered, concentrated to remove molecules <100 kDa, and ultracentrifuged to pellet the EVs (Fig. 1a). To remove non-membranous proteins, protein aggregates, and denatured EVs, a 10-40% Optiprep-based density gradient centrifugation step was performed on the crude EV preparations. Consecutive (top to bottom) Optiprep fractions (10 μl) were subjected to SDS-PAGE. As shown in Fig. 1b, little silver-stained material was recovered from fractions 1 and 2. Samples with similar protein banding patterns (fractions 3-8 and 9-11) were pooled, diafiltered to remove the Optiprep medium, and examined by TEM. Notably, EVs were only observed in the pooled sample from fractions 3-8 (Fig. 1c), but not from fractions 9-11 (Fig. 1d). These results indicated that EVs are produced by USA300 strain and were mainly distributed in fractions containing 20%-35% Optiprep.

**Fig. 1.**
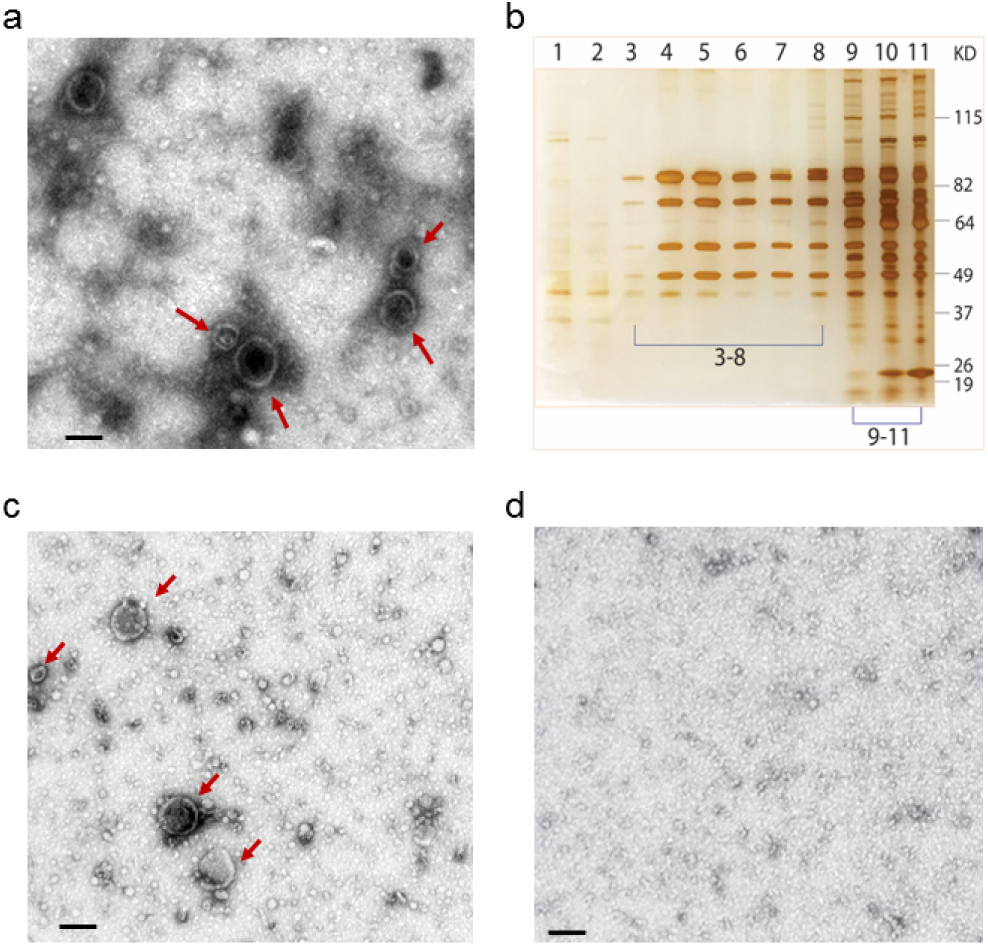
*S. aureus* JE2 produces EVs that can be recovered from culture supernatants after filtration and concentration to remove molecules <100 kDa. **a)** Crude EVs pelleted by ultracentrifugation from a JE2 culture supernatant were imaged by TEM. EVs are shown by red arrows. **b)** EVs were further purified by density gradient ultracentrifugation (Optiprep), and fractions were visualized by silver-stained SDS-PAGE. **c)** Fractions 3-8 were pooled; OptiPrep was removed by diafiltration, and the samples were imaged by TEM. **d)** EVs were not visualized in fractions 9-11. Scale bar: 100 nm.

### Identification of EV-associated proteins by liquid chromatography-tandem mass spectrometry (LC-MS/MS)

To identify the proteins comprising EVs from *S. aureus* JE2, we performed a proteomic analysis of Optiprep-purified EVs using LC-MS/MS. A total of 165 proteins were identified in JE2 EVs (Table S1), including many proteins that are characterized as virulence factors, such as alpha toxin (Hla), leukocidin subunits (LukS-PV, LukF-PV, LukE, LukD, HlgB, and HlgC), surface adhesins (ClfA, ClfB, SdrD, SdrE, Efb, and Ebh), MntC, proteases, and immune evasion factors (Sbi, phenol soluble modulins, catalase, CHIPS, and SodA). Other proteins of interest included penicillin binding proteins, autolysins (Atl, Sle and other putative autolysins predicted to have N-acetylmuramoyl-L-alanine amidase activity), and multiple proteins involved in iron acquisition. A number of lipoproteins with and without characterized functions were also identified in JE2 EVs. Bioinformatic analyses revealed that 76 cytoplasmic proteins were enriched in *S. aureus* EVs, and these represented 46% of all EV proteins. Cell wall associated proteins (n=16), extracellular proteins (n=27), membrane proteins (n=19), and proteins with an unknown localization (n=27) accounted for 10%, 16%, 12%, 16% of total proteins, respectively.

### PSM peptides are involved in the generation of *S. aureus* EVs

The mechanisms underlying EV production by *S. aureus* and other Gram-positive bacteria remain unclear. PSMs are a family of amphipathic, alpha-helical, surfactant-like peptides secreted by *S. aureus*, which are proinflammatory and show cytolytic activity against neutrophils^34,35^. Alpha-type PSMs are required for mobilizing lipoproteins from the staphylococcal cytoplasmic membrane, a process essential for activating TLR2^36^, as well as the export of cytoplasmic proteins, consistent with the membrane-damaging activity of PSMs^5^. Because the cargo of *S. aureus* EVs is enriched for both lipoproteins and cytoplasmic proteins, we evaluated whether PSM peptides were critical for EV generation.

We measured EV production by the WT USA300 LAC strain (the parent strain of JE2), as well as LAC Δ*psmα*, Δ*psmβ*, and Δ*psmαΔpsmβ* mutants^34^. First, we evaluated EV production by dot immunoblot analysis with detection by serum from mice immunized with WT JE2 EVs. Only mutation of *psmα* genes showed a reduced signal in immunoblotting assay for EV production (Fig. 2a). To further substantiate our result, we measured EV yield and particle number by assays for protein content and nanoparticle tracking, respectively. Consistently, mutation of the *psmα* genes significantly reduced *S. aureus* EV yield (Fig. 2b) and particle number (Fig. 2c). The Δ*psmα* and Δ*psmα*Δ*psmβ* double mutant produced comparable levels of EVs when tested with EV yield analysis (Fig. 2b), indicating that psmα peptides play the dominant role in this phenotype. Complemention with pΔTX expressing PSMα1-4 genes, but not the pΔTX vector alone, restored EV production to the Δ*psmα* mutant (Figs. 2b and 2c). Mutation of the *psmα* genes significantly reduced *S. aureus* EV size (Fig. 2c and 2d), whereas the Δ*psmβ* mutant produced EVs of intermediate size compared to that of WT LAC.

**Fig. 2.**
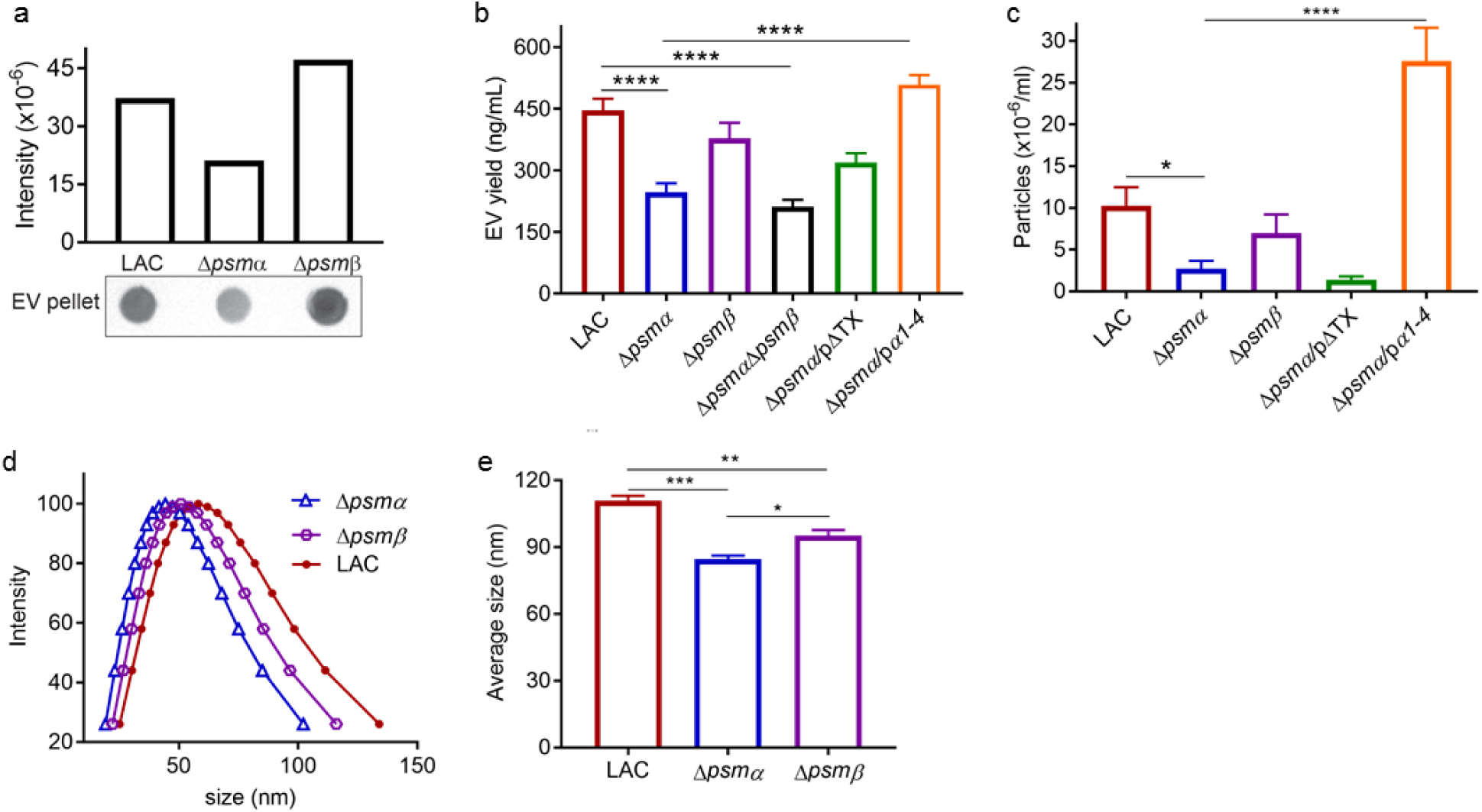
PSMα peptides promote *S. aureus* EV production. **a)** EV production from strain LAC and its isogenic Δ*psmα* and Δ*psmβ* mutants was evaluated by dot-blotting EV suspensions purified from the same volume of bacterial culture, **b)** by quantification of total EV protein abundance, or **c)** by direct EV quantification of EV particles with nanoparticle tracking analysis. Dot-blotting was repeated at least twice, and a representitative result is presented. Signal intensity quantified by Image Studio Lite software is shown above the blot. EV quantification by other methods was calculated from at least three independent experiments and expressed as mean ± SEM. **d)** The size distribution and e) average size of EVs purified from WT and Δ*psmα* and Δ*psmβ* mutants were measured by dynamic light scattering. Data were analyzed using One-way ANOVA with Bonferroni’s multiple comparison test (Fig. 3b and 3c) or with Tukey’s multiple comparison test (Fig. 3e). * P<0.05, ** P<0.01, *** P<0.001, **** P<0.0001.

### PGN cross-linking modulates EVs production

Unlike OMVs produced by Gram-negative microbes, *S. aureus* cytoplasmic membrane-derived EVs must traverse a PGN cell wall structure before cellular release. To determine whether the degree of PGN crosslinking affected *S. aureus* EV biogenesis, we cultured *S. aureus* JE2 in medium with a sublethal concentration (0.2 μg/ml) of penicillin G (PenG) that has been shown to decrease PGN cross-linking^37^. Treatment with a sublethal concentration (0.1 μg/ml) of Em served as antibiotic control that has no effect on PGN cross-linking. Compared to EVs recovered from untreated cultures or cultures incubated with Em, the EV yield from PenG-treated cultures was distinctly higher (Fig. 3a). When the EV protein content was quantified from a fixed volume of culture left untreated or treated with sublethal antibiotic concentrations, we observed a 10-fold increase in EV yield from PenG-treated cultures (Fig. 3b).

**Fig. 3.**
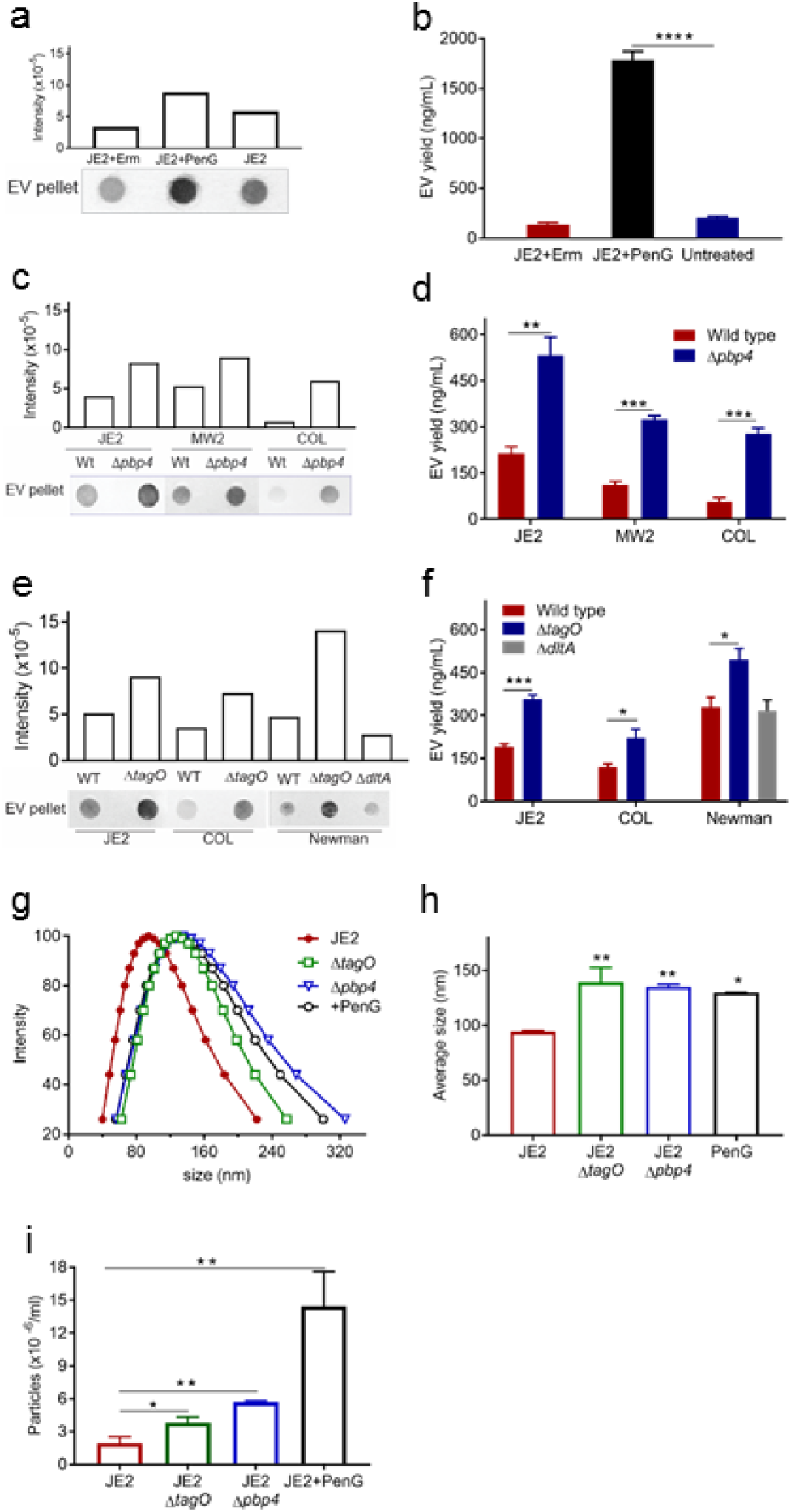
Reductions in PGN crosslinking and lack of WTA increased the production and size of *S. aureus* EVs. **a)** Dot blots were performed on JE2 EVs treated with subinhibitory concentrations of PenG or Em and probed with mouse EV antiserum. **b)** EV protein abundance was quantified and expressed as nanograms EV protein per ml culture. **c)** EV production from *S. aureus* strains JE2, MW2, COL, or their isogenic Δ*pbp4* mutants was evaluated by dot-blotting EV suspensions or **d)** by quantification of total EV protein yield. **e)** EV production from strains JE2, COL, Newman, and their Δ*tagO* and Δ*dltA* mutants was evaluated by dot-blotting EV suspensions or **f)** by quantification of total protein yield. **g)** The size distribution and **h)** average size of EVs isolated from JE2, PenG treated JE2, and Δ*pbp4* and Δ*tagO* mutants were measured by dynamic light scattering. **i)** EV particles from JE2, PenG treated JE2, and Δ*pbp4* and Δ*tagO* mutants were quantified by nanoparticle tracking analysis. The dot immunoblot assay was repeated at least twice with similar results; a representative blot is shown. EV protein yield and EV particle quantification experiments were calculated from at least three independent experiments and expressed as mean ± SEM. The data were analyzed using one-way ANOVA with Dunnett’s multiple comparison test (Fig. 3b, 3h, and 3i) or using Student’s t-test (Fig. 3d and 3f). * P<0.05, ** P<0.01, *** P<0.001, **** P<0.0001.

S. *aureus* penicillin binding protein 4 (PBP4) is a carboxypeptidase that is essential for secondary crosslinking of PGN, and a *pbp4* mutant shows a significant reduction in PGN crosslinking^37,38^.As predicted, both dot blot (Fig. 3c) and EV protein yield assays (Fig. 3d) showed increased EV production by JE2Δ*pbp4*, and the protein yield was 3-fold higher than the wild-type JE2 strain. We also measured EV production in MRSA isolates MW2, COL, and their Δ*pbp4* mutants; the relative increase in EV yield in the mutant strains (Fig. 3c and 3d) was consistent with that of JE2 Δ*pbp4*.

WTA is a PGN-anchored glycopolymer that is major component of the *S. aureus* cell wall and plays a critical role in cell wall homeostasis^2^. The *tagO* gene encodes an *N*-acetyl glucosamine-phosphate transferase enzyme that catalyzes the first step in WTA biosynthesis^39,40^, and deletion of *tagO* gene abrogates *S. aureus* WTA production^41^. Compared to the WT strains JE2, COL, and Newman, *tagO* mutants showed an enhanced signal in the dot immunoblot assay for EV production (Fig. 3e). Likewise, quantitative analysis of EV protein yield showed that all three *tagO* mutants produced significantly more EVs than the parental isolates (Fig. 3f). Thus, WTAs negatively modulate *S. aureus* EV production, consistent with reports showing that *tagO* mutants are characterized by diminished PGN cross-linking^42^. The WTA backbone is decorated with ester-linked D-ala residues, which confer a zwitterionic charge to the polymer^2,43^. As shown in panels e and f of Fig. 3, production and yield of EVs by the *ΔdltA* mutant were similar to that of the parental strain Newman.

To determine whether EV size was affected by reduced PGN crosslinking, the size distribution of purified EVs was measured by dynamic light scattering (DSL). Treatment of JE2 cultures with 0.2 μg/ml PenG or mutation of *pbp4* or *tagO* resulted in a significant increase in the size distribution of EVs (Fig. 3g), as well as an increased EV average size (Fig. 3h) compared to untreated WT EVs. Because enhanced EV production and yield associated with reduced PGN crosslinking might be a result of larger EVs that would carry an increased cargo load, we quantified EVs by nanoparticle tracking analysis. As shown in Fig. 3i, treatment of JE2 cultures with 0.2 μg/ml PenG or mutation of *pbp4* or *tagO* resulted in suspensions containing significantly greater numbers of EV particles per ml compared to untreated WT EVs. Taken together, our data indicate that *S. aureus* EV production is inversely proportional to the degree of PGN crosslinking.

### Autolysin Sle1 modulates the release of EVs

Atl and Sle1 belong to a family of PGN hydrolases that plays a critical role in separation of daughter cells^44,45^. In addition, Atl has been shown to modulate the excretion of a subset of staphylococcal cytoplasmic proteins^4^. The staphylococcal autolysins Atl and Sle1 are abundant proteins in JE2 EVs (Table S1), as well as in *S. aureus* EV preparations characterized previously^16^.

To determine whether PGN-hydrolases facilitate the release of EVs by altering the thick cell wall of Gram positive bacteria, we compared EV production from *atl* and *sle1* mutants with that of strains JE2 and Newman. Although both mutants showed reduced EV production (Fig. 4a), the reduction in yield was only significant in the *sle1* mutants (Fig. 4b). Similar results were obtained by nanoparticle tracking analysis wherein EV purification from the *sle1* mutant, but not the *atl* mutant, yielded a significantly lower EV concentration compared with that of the WT strain JE2 (Fig. 4c). To determine whether autolysin activity affected the size distribution of EVs, we evaluated purified EVs by dynamic light scattering. As shown in Fig. 4d and 4e, both *atl* and *sle1* mutations exhibited a reduced EV size compared to WT JE2 EVs.

**Fig. 4.**
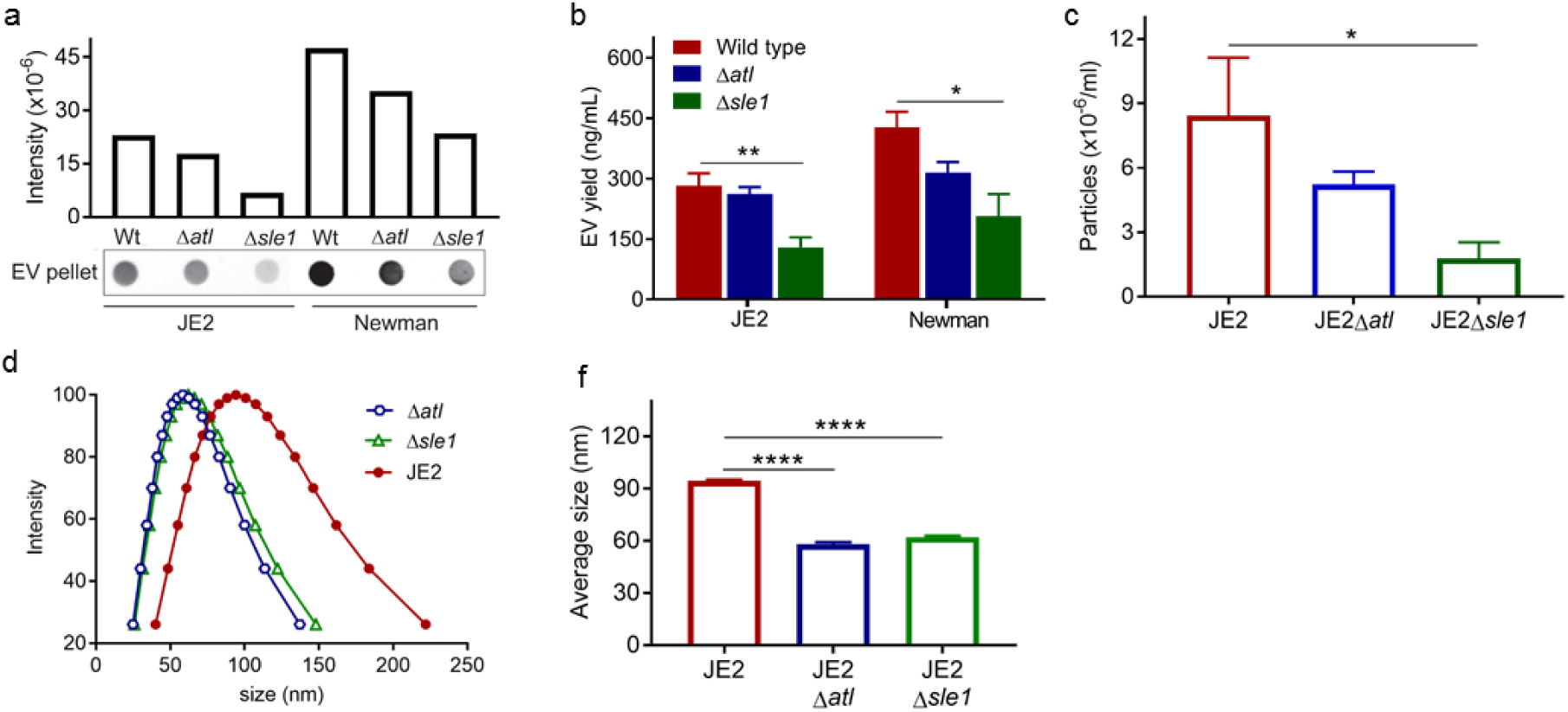
The autolysin Sle1, but not Atl, promoted EV biogenesis. **a)** EV production from JE2 and Δ*atl* and Δ*sle1* mutants was evaluated by dot-blotting EV suspensions from filter-sterilized culture filtrates, **b)** by quantification of total EV protein abundance, or **c** by EV quantification using nanoparticle tracking analysis. **d)** The size distribution and **e)** average size of EVs isolated from JE2 and the Δ*atl* and Δ*sle1* mutants were measured by dynamic light scattering. The dot immunoblot assay was repeated at least twice with similar results; a representative blot is shown. EV protein yield and EV particle quantification experiments were calculated from at least three independent experiments and expressed as mean ± SEM. The data were analyzed using oneway ANOVA with Dunnett’s multiple comparison test (Fig. 4b, 4c, and 4e), For all panels, * P<0.05, ** P<0.01, *** P<0.001, **** P<0.0001.

### The influence of capsular polysaccharide (CP) on *S. aureus* EV production

Streptococcal CP production was shown to hinder EV release by *Streptococcus pneumoniae*^7^ but not *S. pyogenes*^8^. To determine whether the presence of *S. aureus* CP impacted EV biogenesis, we evaluated EV production by a number of isogenic CP+ and CP-strains. The EV pellet was derived by ultracentrifugation of filter-sterilized culture supernatants and would not contain soluble CP. As shown in Fig. 5a, the CP phenotype had no obvious impact on the EV dot blot signal derived from WT or CP-mutants of strains Newman or 6850. Similarly, when we complemented the genetic defect in the *cap5* locus of USA300 strain 923 with pCap17, the strain produced CP5^46,47^, but there was no effect on the EV signal levels achieved by dot blotting (Fig. 5a). Likewise, CP+ and isogenic CP-strains of Newman, 6850, and 923 produced comparable protein yields of EVs (Fig. 5b). Thus, *S. aureus* CP did not influence the yield or block the release of EVs from *S. aureus* Newman (CP5+), 6850 (CP8+), or a CP5+ USA300 isolate.

**Fig. 5.**
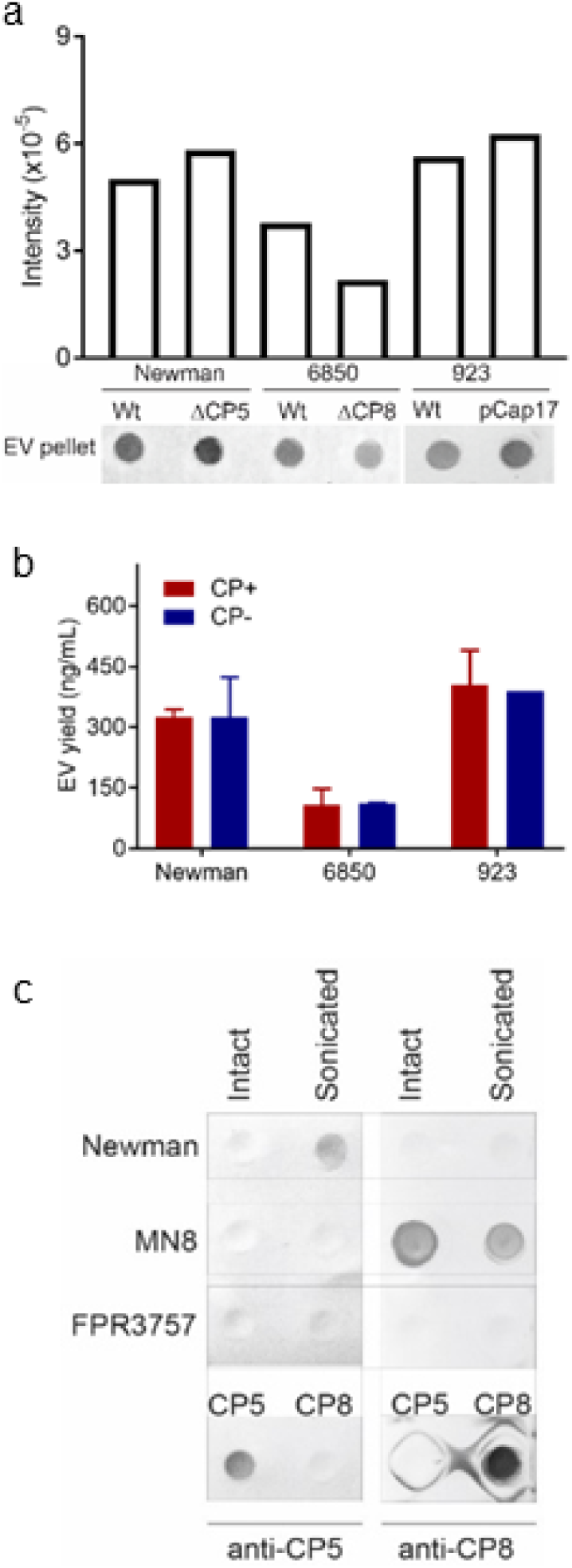
*S. aureus* CP is associated with EVs but does not impact EVs biogenesis. **a)** EV production from encapsulated *S. aureus* (Newman, 6850, 923 [pCP5]) or their CP-negative counterparts (NewmanΔ*cap5O*, 6850Δ*capHIJK*, and 923) was evaluated by dot-blotting EV suspensions or **b)** by quantification of total EV protein yield. **c)** CP5 or CP8 was detected in intact or sonicated EVs (35 μg) from strains Newman, MN8, or FPR3757 by immunoblots probed with 1.2 μg/ml CP5-specific mAb 4C2 or CP8-specific mAb 5A6. Controls included purified CP5 (15 μg) and CP8 (15 μg). The dot immunoblot assay was repeated at least twice with similar results; a representative blot is shown. EV protein yield were calculated from at least three independent experiments and expressed as mean ± SEM.

To investigate whether CP antigens were associated with *S. aureus* EVs, we performed CP immunoblots on EVs prepared from strains Newman (CP5+), MN8 (CP8+), and USA300 FPR3757 (CP-). We tested both intact and sonicated EVS since CP antibodies would react with surface-associated CP antigens on intact EVs, whereas intravesicular CP antigens would only be detected in sonicated EV preparations. Figure 5c shows that CP5 was only detected in sonicated, but not intact Newman EVs, whereas CP8 was detected in both intact and sonicated MN8 EVs. Because WT FPR3757 produces no CP, EVs from this strain reacted with neither CP antibody. These data indicate both CP5 and CP8 were associated with EVs produced by CP+ *S. aureus* isolates, although only CP8 was surface exposed. Additional studies outside the scope of this report are needed to confirm and evaluate the prevalence, significance, and mechanism by which CP is surface exposed on EVs prepared from a variety of *S. aureus* isolates.

### Detoxified EVs secreted by a *S. aureus* mutant are immunogenic and represent a novel multicomponent vaccine platform

JE2 EVs packaged multiple antigens including lipoproteins, cytolytic toxins, surface proteins, and enzymes (Table S1). Thus, JE2 EVs could serve as a multivalent *S. aureus* vaccine candidate if the toxicity of the EVs were eliminated. We detoxified *S. aureus* EVs by genetically repressing the expression of cytolytic toxins by mutation of *agr*, an *S. aureus* global regulator. We subsequently deleted *spa* (the gene encoding protein A) in the EV host strain since an *agr* mutant overexpresses Spa, which binds to the F_cy_ domain of immunoglobulin and dampens antibody development by cross-linking the Fab domain of V_H_3-type B cell receptors, resulting in apoptotic collapse of these cells^48^. The JE2Δ*agr*Δ*spa* double mutant served as our *S. aureus* EV vaccine producing host strain. Consistent with previous reports, we demonstrated that the JE2 *agr* mutation significantly inhibited mRNA expression of *hla* and the genes encoding all nine leukocidin subunits (Fig. S1a). EVs from WT JE2, but not the JE2Δ*agr*Δ*spa* mutant, contained native Hla as assessed by western blotting (Fig. S1b). Similarly, by using an antibody reactive with both LukS-PV and LukE subunits, we showed that only EVs from WT JE2 had detectable leukocidin reactivity (Fig. S1b). To further validate our results, we analyzed the protein content of EVs purified from JE2Δ*agr*Δ*spa* by MS. Notably, many of the extracellular proteins that were present in JE2 WT EVs were not detectable in EVs from JE2AagrAspa. However, some antigens such as MntC and FhuD2 that have been shown to protect mice against experimental *S. aureus* infections^49-52^ were still present in EVs from the mutant strain. Neither protein A nor the cytotoxins Hla, Luk-PVL, LukED, HlgCB, SelX, and PSMs were detectable by LC-MS/MS in EVs purified from the JE2Δ*agr*Δ*spa* mutant (Table S2). Although LukAB was still present in EVs from JE2Δ*agr*Δ*spa*, there was ≥86% reduction in the number of peptides detected in the mutant strain (Table S1 and Table S2). Moreover, as indicated below, EVs recovered from the mutant strain showed no residual toxicity toward human leukocytes.

To investigate whether the detoxified JE2 EVs were immunogenic and protective against infection, we immunized mice with 5 μg EVs from JE2Δ*agr* or JE2Δ*agr*Δ*spa* mutants; control mice were given PBS. EVs from either mutant elicited a serum antibody response against sonicated WT EVs, although the antibody level elicited by Δ*agr* EVs was higher than that elicited by Δ*agr*Δ*spa* EVs (Fig. S2a). To examine the antigen profiles from EVs that elicited antibody responses after immunization, a bacterial lysate from the USA300 FPR3757 strain was subjected to SDS-PAGE and immunoblotted with pooled sera from mice immunized with either Δ*agr* EVs or Δ*agr*Δ*spa* EVs. Notably, sera from Δ*agr*Δ*spa* EVs-immunized mice reacted with more bacterial antigens than sera from Δ*agr* EVs-immunized mice (Fig. S2b), suggesting that Δ*agr*Δ*spa* EVs elicited a greater diversity of antibodies than Δ*agr* EVs. To further evaluate the protective efficacy of EVs, the immunized mice were challenged with WT USA300 strain FPR3757. Immunization of mice with EVs from JE2Δ*agr*Δ*spa*, but not EVs from JE2Δ*agr*, provided significant protection against lethal sepsis (Fig. S2c). Preliminary studies indicated that immunization with higher doses of EVs mixed with alum did not enhance immunogenicity (Fig. S2d).

### *S. aureus* engineered EVs (eng-EVs) elicit neutralizing antibodies and protect against lethal sepsis

An ideal multicomponent *S. aureus* vaccine should elicit cytotoxin neutralizing antibodies. Hla is a major secreted staphylococcal cytotoxin, and its production has been associated with severe infections caused by community-acquired MRSA^53^. Immunization against a nonpore-forming Hla variant (Hla_H35L_) prevents experimental *S. aureus* pneumonia, skin abscesses, and lethal peritonitis^54-56^. To enhance the protective efficacy of detoxified EVs from JE2Δ*agr*Δ*spa*, we engineered JE2 to package nontoxic Hla_H35L_^57^ and the LukE monomer within eng-EVs. LukED is a member of the *S. aureus* family of bicomponent leukotoxins and is detected in 82% of blood isolates and 61% of nasal isolates^58^. LukED targets both human and murine neutrophils, macrophages, T cells, dendritic cells, NK cells, and erythrocytes^59,60.^

We expressed nontoxic Hla_H35L_ and LukE in strain JE2Δ*agr*Δ*spa* under control of the *spa* promoter. Because the activity of the *spa* promoter is enhanced in an Δ*agr* genetic background, the mRNA levels of Hla_H35L_ and LukE expressed in JE2Δ*agr*Δ*spa* were dramatically increased compared to expression in JE2Δ*agr*Δ*spa* or JE2Δ*agr*Δ*spa* with the empty vector (Fig. S1c). As predicted, both Hla_H35L_ and LukE were detected by Western blot in engineered EVs (eng-EVs) isolated from recombinant strain JE2Δ*agr*Δ*spa* (pHla_H35L_-LukE) (Fig. S1b).

To evaluate the relative toxicity of EVs prepared from WT strain JE2 and JE2Δ*agr*Δ*spa* vs. eng-EVs from JE2Δ*agr*Δ*spa* (pHla_H35L_-LukE), we incubated the EVs in vitro with three different cell types. A549 cells are susceptible to Hla-mediated cytolysis, and WT strain JE2 EVs were toxic at concentrations as low as 1 μg/ml. In contrast, JE2Δ*agr*Δ*spa* mutant EVs and the eng-EVs from JE2Δ*agr*Δ*spa* (pHla_H35L_-LukE) exhibited negligible toxicity (Fig. S3a). HL60 cells are resistant to Hla-mediated lysis, but they are susceptible to the cytolytic activity of the *S. aureus* leukocidins (including HlgAB, HlgCB, PVL-SF, LukED, LukAB, and phenol soluble modulins [PSMs]). EVs isolated from strain JE2, but not the Δ*agr*Δ*spa* mutant or eng-EVs, were cytolytic for HL60 cells at concentrations as low as 1 μg/ml (Fig. S3b). Rabbit erythrocytes are susceptible to Hla, PSMs, and the leukocidins HlgAB and LukED^61,62^. EVs isolated from WT strain JE2 exhibited significant hemolytic activity at concentrations as low as 1 μg/ml, but no hemolytic activity resulted from EVs prepared from the Δ*agr*Δ*spa* mutant or eng-EVs, even at 20 μg/ml (Fig. S3c). These data demonstrate that the eng-EVs were nontoxic in vitro for mammalian cells.

We immunized mice on days 0. 14, and 28 with 5 μg EVs from JE2Δ*agr* or JE2Δ*agr*Δ*spa* mutants; control mice were given 5 ug bovine serum albumin (BSA). Whereas sera from mice immunized with both eng-EVs and Δ*agr*Δ*spa* EVs, but not BSA, reacted by ELISA with sonicated WT JE2 EVs (Fig. 6a), only mice given the eng-EVs responded with antibodies to purified Hla (Fig. 6b) or LukE (Fig. 6c). These data indicate that recombinant proteins packaged within *S. aureus* EV are immunogenic.

**Fig. 6.**
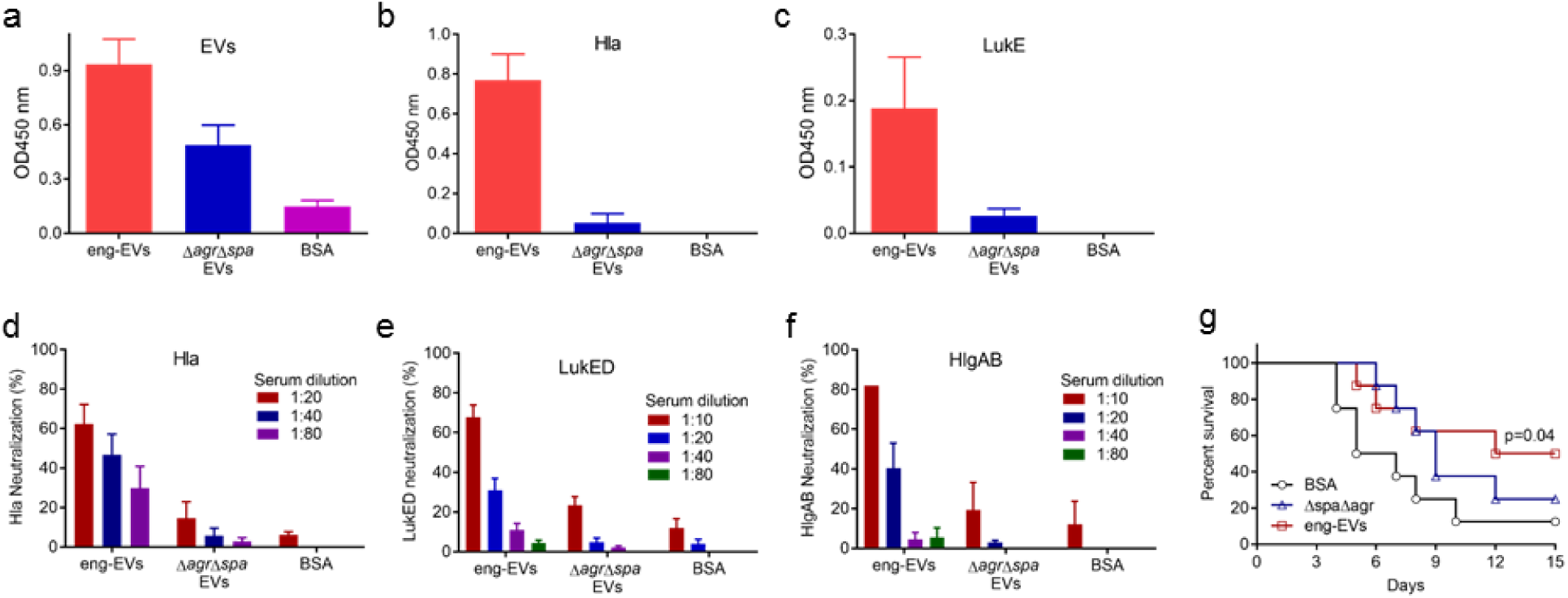
Immunogenicity and protective efficacy in mice of eng-EVs. Antibody levels in sera (diluted 1.100) from mice immunized with eng-EVs were analyzed on ELISA plates coated with **a)** JE2 sonicated EVs, **b)** Hla, or **c)** LukE. Data were expressed as mean ± SEM. The neutralizing activity of sera from mice immunized with BSA or different EV preparations was determined by either incubating serial dilutions of sera with **d)** Hla, **e)** LukED, or **f)**, or HlgAB for 1 h at 37°C before adding target cells. Control cells were incubated with toxins but no sera. Data are expressed as percent neutralization ± SEM. Mice (n=8) immunized with different JE2 EV preparations were challenged IV with 2 × 10^8^ CFU *S. aureus* NRS685 **(g)**. Survival (comparing EV-immunized mice vs. BSA immunized mice) was analyzed with the log rank test.

To examine whether the antibodies elicited in mice by the eng-EV vaccine were functional, toxin neutralizing assays were performed. Sera from mice immunized with eng-EVs effectively neutralized Hla at dilutions ranging from 1:20 to 1:80 (Fig. 6d). In contrast, neutralizing antibodies were low or undetectable in serum from mice given BSA or Δ*agr*Δ*spa* EVs. Similarly, sera from mice immunized with eng-EVs, but not BSA or Δ*agr*Δ*spa* EVs, were able to effectively neutralize LukED at dilutions ranging from 1:10 to 1:20 (Fig. 6e). Sera from mice immunized with eng-EVs also neutralized leukocidin HlgAB (Fig. 6f), but not PVL-SF or HlgCB leukotoxins.

The immunized mice were challenged with USA500 strain NRS685, a PVL-negative MRSA bacteremia isolate. We chose this strain because the PVL-S and PVL-F subunits can interact with LukE and LukD to form inactive hybrid complexes, and this influences LukED-mediated *S. aureus* virulence in mice^63^. As shown in Fig. 6g, immunization with eng-EVs, but not Δ*agr*Δ*spa* EVs, protected 50% of the mice in the lethal sepsis model.

## Discussion

Membrane vesicles, produced by mammalian cells, fungi, and bacteria, is an evolutionarily conserved secretory pathway that allows cell-free intercellular communication^64-66^. Microbial EVs encapsulate cargo that include lipids, proteins, glycans, and nucleic acids, which have been shown to play roles in microbial physiology, pathogenesis, and the transmission of biological signals into host cells to modulate biological processes and host innate immune responses^64,65,67,68^. In Gram-negative bacteria, EVs are generated by pinching off the outer membrane, but the mechanism(s) by which EVs escape the thick cell walls of Gram-positive bacteria, mycobacteria, and fungi is unknow. Once shed, *S. aureus* EVs can undergo cholesterol-dependent fusion with host cell membranes to deliver their toxic cargo^28^. *S. aureus* EVs have been shown to be produced in vivo during experimental pneumonia in mice^28^. In this report, we demonstrate unique properties associated with EV production by JE2, a *S. aureus* USA300 strain that is representative of the CA-MRSA clone that has rapidly disseminated in the United States. Similar to EVs characterized from other *S. aureus* isolates^16,17,28^, JE2 EVs encapsulate an array of bacterial antigens, including lipoproteins, exotoxins, and cytoplasmic proteins.

In an effort to better understand the multiple stages of EV biogenesis in *S. aureus*, we evaluated putative factors that modulate the membrane and PGN related steps of EV release. We first explored the influence of PSMs on the membrane step of this process. PSMs are a group of small alpha helical peptides that have surfactant-like properties and potent cytolytic activity for leukocytes and other host cells, as well as membrane disturbing activity on the producing *S. aureus* cell^5^. PSMα peptides are 20-22 amino acids in length, whereas PSMβ peptides are 43-45 amino acids in length^35^. In our studies, PSMα peptides, but not PSMβ peptides, supported the generation of EVs from *S. aureus*. EVs from the PSMα mutant were less abundant and smaller in size compared with WT EVs. Although PSMβ peptides did not significantly reduce EV yield, the smaller size of the mutant EVs suggest that PSMβ peptides may play an accessory role in EV biogenesis. In an *S. aureus* mutant that lacks the PSM transporter protein, PSMs accumulate intracellularly, causing cytoplasmic membrane perturbations^69^. A recent study reported that PSMα peptides induced the release of cytoplasmic proteins, lipids, nucleic acids, and ATP into *S. aureus* culture supernatants, and that this effect was mediated by the membrane-damaging activity of PSMα^5^. Surfactants or surfactant-like proteins with amphipathic helical structures have been shown to insert into lipid monolayers and generate local deformatio^70,71^. PSMs, due to their surfactant-like activity as well as amphipathic helical structure, may enhance membrane curvature under cytoplasmic turgor pressure, resulting in membrane disruption and the formation of EVs.

The *S. aureus* cell envelope is comprised of a thick, highly cross-linked PGN layer, proteins, and glycopolymers like WTA and CP. When we assessed the role of the PGN layer on EV release, we found that highly crosslinked PGN serves as a barrier for EV biogenesis. Treatment of *S. aureus* with sublethal concentration of penicillin G or genetic inactivation of *pbp4* or *tagO*, which result in reductions in PGN cross-linking, resulted in a significant increase in EV production, as well as the average size of released EVs. This inverse correlation between PGN cross-linking and EV yield was not unique to USA300 strain JE2 but was also observed with *S. aureus* strains MW2, COL, and Newman. WTA has been shown to be critical for PGN-crosslinking by regulating PBP4 localization to the septation site^42^. A secondary mechanism by which WTA regulates EV production is via its ability to control the activity of Atl and Sle1 - not only by preventing their binding to *S. aureus* cell wall PGN^72^ ^73^, but also by creating an acidic milieu that limits Atl PGN hydrolase activity^74^. Consequently, autolytic activity is not localized to the septum area in a *tagO* mutant but is spread throughout the cell surface, likely facilitating EV release. Of note, Schlag et al. reported that a *tagO* mutant showed an altered cell surface with bobble‐ and hairy-like protrusions^72^, which may represent EVs. Although we do not yet fully understand the mechanism(s) of EV generation in Gram positive bacteria, it seems logical that a poorly cross-linked cell wall or a cell wall lacking WTA would lessen the barrier to EV release and generate larger EVs as a result of larger pores within the PGN structure.

Autolysins that cleave the PGN barrier also impact the biogenesis of *S. aureus* EVs. Atl and Sle1 localize to the septum during cell division where they exhibit peptidoglycan hydrolase activity, resulting in separation of the daughter cells^72^ ^73^. Sle1 is a 32 kDa protein comprised of an N terminal cell wall binding domain and a C terminal catalytic domain with N-acetyl muramyl-L-alanine amidase activity. In contrast, Atl is a 138 kDa bifunctional PGN hydrolase that is processed to yield a 62 kDa protein with amidase activity (similar to that of Sle1) and a 51 kDa protein with endo-β-N-acetyl glucosaminidase activity. In addition to its activity in cell separation, Atl is also involved in cell wall turnover and penicillin- or detergent-induced bacterial autolysis. Both Atl and Sle1 proteins are present in EVs isolated from USA300 JE2, as well as ATCC 14458^16^, although Atl is more abundant. Nonetheless, deletion of Sle1, but not Atl, significantly reduced *S. aureus* EV production. Pasztor et al. reported that an SA113 *atl* mutant overexpressed eight putative secondary PGN hydrolases both at the transcriptional and at the protein levels, highlighting the supplementary role of these alternative autolysins in the absence of Atl^4^. This observation may at least partially explain why JE2Δ*atl* and NewmanΔ*atl* showed only a modest reduction in EV yield. Mutations in either *sle1* or *atl* resulted in a significant decrease in EV size. Although both autolysin activities are localized to the *S. aureus* septum region, EVs have been visualized surrounding the bacterial surface^7,16,75^. A recent report demonstrated differential roles for Atl and Sle1 during cell division and separation^76^. Whereas Sle1 could be visualized over the entire septal surface, Atl localized only at the external (surface-exposed) edge of the septum^77^. How autolysins modulate EV release from the cell wall or whether this process is spatially or temporally regulated remains to be determined.

*S. aureus* CPs were associated with or packaged within EVs isolated from *S. aureus* strain Newman and MN8 in our study. We reported that *S. aureus* CP was shed from broth-grown *S. aureus* cells^78^, and it is feasible that EVs could serve as a vehicle to liberate CP from the cell envelope. The *S. pneumonia* capsule was reported to hinder EV release in this pathogen^7^, whereas no effect was observed on EV yield in strains with or without the hyaluronic capsule of *S. pyogenes^8^*. Whether these streptococcal CPs are present as EV cargo in these pathogens was not addressed. In our hands, *S. aureus* CPs did not hinder the release of EVs from encapsulated *S. aureus*. Although EV yield varied among different isolates, we recovered similar quantities of EVs from isogenic strains that varied only in CP production. The glucuronoxylomannan capsule of *Cryptococcus neoformans* has been identified as a component of EVs from this fungal pathogen^79^, and polysaccharide A from *Bacteroides fragilis* was shown to be packaged into OMVs that were capable of inducing immunomodulatory signaling in dendritic cells^68^. Ongoing studies in our laboratory will address whether *S. aureus* EV-host cell interactions impact of the pathogenesis of staphylococcal disease.

Extensive efforts have been made to develop vaccines against *S. aureus* infections in humans. Although a vaccine to prevent *S. aureus* disease is still not available, a growing body of evidence has suggested that a successful vaccine should target multiple antigens (toxoids, adhesins, and anti-phagocytic polysaccharides) that play distinct roles in *S. aureus* pathogenesis. Immunization with *S. pneumoniae* EVs protected mice against lethality^7^. Moreover, immunization of mice with native EVs from *S. aureus* ATCC 14458 elicited a robust T cell response and protected mice against pneumonia^27^. Although the latter study demonstrated the immunogenicity and potential of staphylococcal EVs as a vaccine platform, limitations of the report include challenge with the homologous strain and failure to acknowledge the documented cytolytic activity of EVs prepared from WT S. *aureus*^28^.

Because strain JE2 produced EVs that were cytotoxic in vitro when incubated with eukaryotic cells, we mutated *agr*, an *S. aureus* global regulator. As predicted, EVs from JE2Δ*agr* showed undetectable toxicity against epithelial cells, neutrophils, and erythrocytes. Moreover, LCMS/MS analyses revealed that the cytolytic toxins were present in EVs from the WT strain but not the Δ*agr*Δ*spa* double mutant. We also deleted *spa* in the EV host strain since an *agr* mutant overexpresses Spa, which binds to immunoglobulins by their Fc fragment and dampens antibody development by cross-linking the Fab domain of V_H_3-type B cell receptors^48^. Indeed, sera from mice immunized with JE2Δ*agr*Δ*spa* EVs reacted by immunoblot with a greater diversity of *S. aureus* proteins than sera from mice vaccinated with JE2Δ*agr* EVs. Moreover, mice immunized with JE2Δ*agr*Δ*spa* EVs showed a significant reduction in lethality provoked by WT USA300 strain FPR3757 compared to mice given PBS or JE2Δ*agr* EVs.

To enhance the protective efficacy of Δ*agr*Δ*spa* EVs as a vaccine platform, nontoxic Hla_H35L_ and LukE were expressed in JE2Δ*agr*Δ*spa* under the control of the agr-derepressed *spa* promoter. Immunization with purified nontoxic Hla_H35L_ prevents lethal pneumonia and lethal peritonitis and reduces the incidence of necrotic skin abscesses^54-56^. *S. aureus* leukocidins comprise a family of pore-forming toxins produced by *S. aureus* that target monocytes, lymphocytes, neutrophils, and macrophages - the very cells responsible for resolution of bacterial infection. These “eng-EVs” elicited antibodies in the sera of immunized mice that reacted with Hla and LukE by ELISA and neutralized the cytolytic activity of Hla, LukED, and HlgAB in vitro.

Mice immunized with eng-EVs, Δ*agr*Δ*spa* EVs, or BSA were challenged with USA500 strain NRS685, a PVL-negative MRSA bacteremia isolate. LukED has been shown to enhance lethality in mice challenged with S. *aureus^80^*, and the presence of PVL modulates LukED-mediated *S. aureus* virulence in mice^63^. Immunization with eng-EVs, but not Δ*agr*Δ*spa* EVs, protected 50% of the mice in the lethal sepsis model. Protective efficacy against additional *S. aureus* strains and in additional infection models remains to be evaluated. Over-expression of additional antigens that have been shown to protect mice against experimental *S. aureus* infections, such as MntC and FhuD2^49-51,81^, in second-generation eng-EVs may yield a more efficacious vaccine. LC-MS/MS analysis of EVs from both WT JE2 and the Δ*agr*Δ*spa* mutant strain contained multiple lipoproteins. As a predominant TLR2 ligand, lipoproteins have been increasingly used as novel adjuvant components^82,83^ because they are potent activators of host innate immunity and can mediate humoral and cell mediated immune responses.

In summary, we have generated, purified, and characterized EVs isolated from *S. aureus* USA300, the predominant CA-MRSA clone in United States. Our study revealed that *S. aureus* PSMs are central for EVs generation by targeting the cytoplasmic membrane. Likewise, the Sle1 autolysin is critical for the release of EVs from *S. aureus* cell wall. Whereas mutations in Atl or CP production did not affect EV yield, PBP4 and WTA promote PGN cross-linking and consequently diminished EV production. Our study elucidates certain mechanisms whereby *S. aureus* produces and sheds EVs (Fig. 7) and will ultimately further our understanding of bacterial physiology and pathogenesis. We designed and created eng-EVs as a novel vaccine platform against *S. aureus* infection. Detoxified EVs that over-produced Hla_H35L_ and LukE were immunogenic, elicited toxin neutralizing antibodies, and protected mice in a *S. aureus* lethal sepsis model, indicating that these naturally produced vesicles have potential as a noval vaccine platform.

**Fig. 7.**
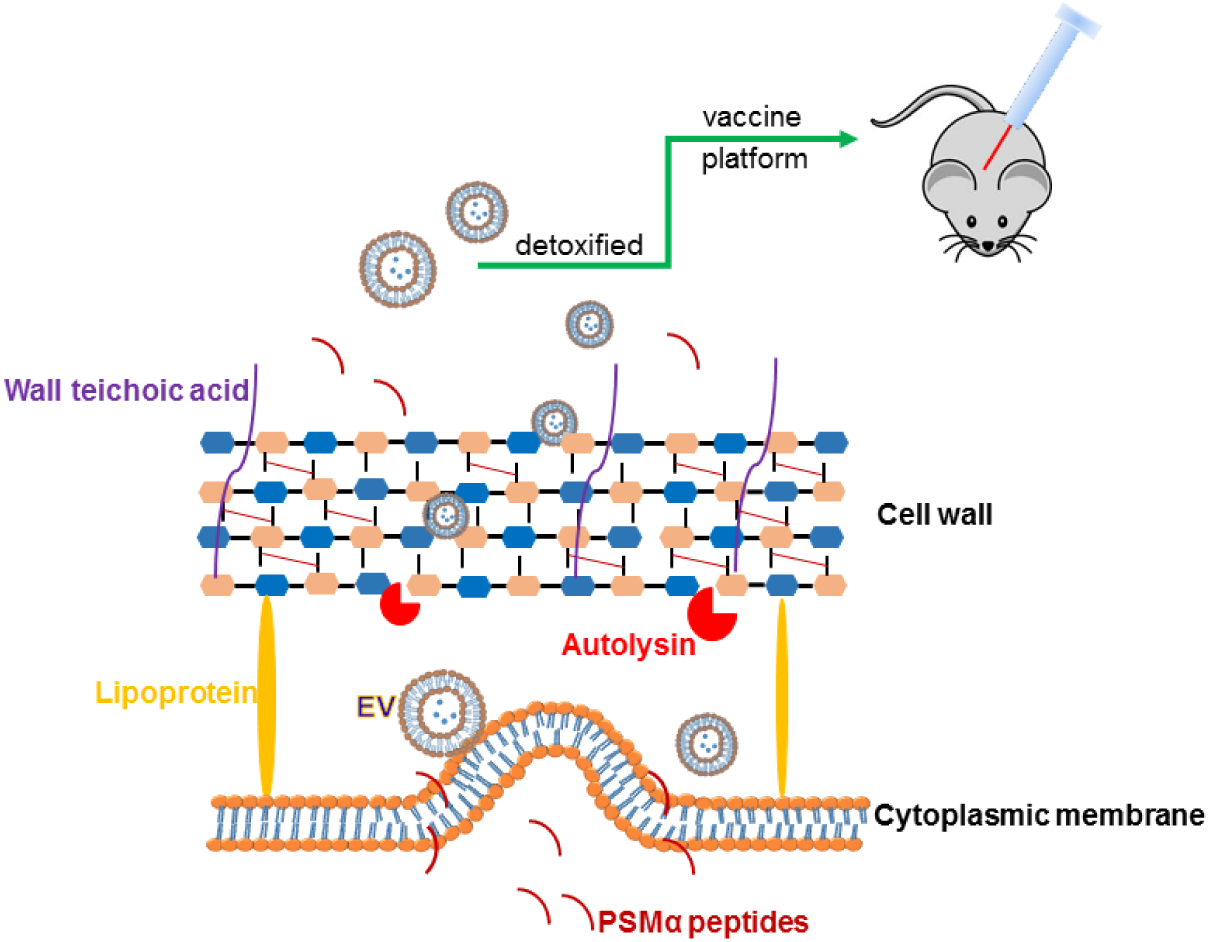
Proposed mechanisms underlying extracellular vesicle (EV) production by *S. aureus*. EVs are generated from the cytoplasmic membrane, and this process is promoted by *S. aureus* PSMα peptides, which have surfactant-like activity, causing membrane disruption. Membrane-derived EVs must also traverse the highly cross-linked *S. aureus* peptidoglycan barrier, and the extent of cell wall cross-linking modulates the efficiency of EV production. Autolysins, such as Sle1, facilitate EV release by hydrolyzing peptidoglycan, particularly at sites of active cell division. We mutated *S. aureus* to render its EVs nontoxic, and then genetically engineered the mutants to package detoxified antigens in EVs. These recombinant EVs were immunogenic in mice and enhanced protective efficacy in a sepsis model of infection.

## Materials and Methods

### Bacterial strains and plasmids

*S. aureus* isolates (listed in Table S3) were cultivated overnight with aeration in tryptic soy broth (TSB; Difco) at 37°C. *Escherichia coli* strain XL-10 (Agilent), used in DNA cloning experiments, was grown at 37°C in Luria Broth (LB; Difco). *S. aureus - E. coli* shuttle vector pCU1^84^ was used for cloning and expression of appropriate genes in *S. aureus*. Antibiotics were added in the following concentrations: penicillin G (penG; 0.2 μg ml^-1^), ampicillin (Amp; 100 μg ml^-1^), erythromycin (Em; 5 μg ml^-1^), chloramphenicol (Cm; 10 μg ml^-1^), kanamycin (Kan; 50 μg ml^-1^), or tetracycline (Tet; 5 μg ml^-1^).

### DNA manipulation

Fey et al. derived *S. aureus* JE2 from the USA300 strain LAC by curing it of plasmids^85^, rendering it sensitive to Em. The *agr* mutation (Δ*agr.tetM*) was transduced from *S. aureus* RN6911^86^ to wild-type (WT) JE2 using bacteriophage φ80α with selection for Tet resistance. To construct the JE2 Δ*agr*Δ*spa* double mutant, the *spa* mutation was transduced from JE2 (*spa*::*ermB*) to JE2Δ*agr* by φ80α transduction. The *pbp4* mutation was transduced from JE2 (Δ*pbp4*::*ermB*) to WT MW2 and COL by φ80α transduction with selection for Em resistance. All mutants were confirmed by PCR using the primers listed in Table S4. ELISA results confirmed the phenotype of the Δ*spa* mutant, and the *agr* mutants lost their hemolytic phenotype. To construct the WTA mutants, the *tagO* mutation was transduced from SA113 Δ*tagO* (pRB*tagO*) to WT JE2 and COL with φ80α with selection for Em resistance. Mutants were confirmed by PCR and acquisition of resistance to lysis by φ80α.

To construct a shuttle vector for expression of Hla_H35L_ and LukE, the *spa* promoter, *hla*_H35L_, and *lukE* genes were amplified from *S. aureus* strains JE2, DU1090 (pHla_H35L_), and FRP3757, respectively. To drive the expression of *hla*_H35L_, its sequence was fused to the 3’ terminus of the *spa* promoter containing the ribosome binding site by overlapping PCR. The *P*_spa^-^_*hla*_*H35L*_ fusion sequence was cloned into the shuttle plasmid pCU1 with restriction enzymes HindIII and SalI. The amplified *lukE* sequence containing a ribosome binding site was inserted into pCU1 with restriction enzymes SalI and EcoRI. The resulting plasmid *pCU1-P*_spa^-^_*hla*_*H35L*_-*lukE* was verified by enzyme digestion and DNA sequencing. To construct JE2Δ*spa*Δ*agr* expressing nontoxic Hla_H35L_ and LukE, *pCU1-P*_spa^-^_*hla*_H35L_-*lukE* was transformed into RN4220 by electroporation and then transduced with φ80α to JE2Δ*spa*Δ*agr*, selecting for Cm resistance.

### Isolation and purification of EVs

Isolation of EVs from *S. aureus* was performed as previously described^7,16^ with minor modifications. *S. aureus* was cultivated in TSB with shaking to an OD_650 nm_ of 1.2. The culture supernatant was filtered and concentrated 25-fold with a 100-kDa tangential flow filtration system (Pall Corp.). The retentate was filtered again before centrifugation at 150,000 g for 3 h at 4°C to pellet the vesicles and leave soluble proteins in the supernatant. The EV pellet was suspended in 40% Optiprep density gradient medium (Sigma) and overlaid with gradient layers of Optiprep ranging from 35% to 10%. After centrifugation at 139,000 g for 16 h at 4°C, 1 ml fractions were removed sequentially from the top of the gradient. Each fraction was subjected to SDS-PAGE and stained with a Thermo Fisher silver staining kit. Fractions with a similar protein profile on SDS-PAGE were pooled, and the Optiprep medium was removed by diafiltration with phosphate-buffered saline (PBS; 10 mM Na_2_HPO_4_, 2 mM KH_2_PO_4_, 2.7 mM KCl, and 137 mM NaCl, pH 7.4) using an Amicon Ultra-50 Centrifugal Filter Unit. The diafiltered retentate was filtered (0.45 μm) and stored at 4°C. EV protein concentrations were determined by using a Protein Assay Dye Reagent (Bio-Rad). The size distribution and diameter of vesicles was measured using a ZetaPALS dynamic light scattering detector (Brookhaven Instruments Corp.). Nanoparticle tracking analysis (NTA) was performed by purifying EVs from 100 ml bacterial cultures, as described above. The number of EV particles recovered from individual cultures (and suspended in 1 ml PBS) was determined using a Nanosight NS300 Sub Micron Particle Imaging System (Malvern), as previously described^87^.

### Electron microscopy of *S. aureus* EVs

Five microliters of *S. aureus* EVs were adsorbed for 1 min to a carbon coated grid that was made hydrophilic by a 30-sec exposure to a glow discharge. The samples were stained with 0.75% uranyl formate for 30 sec and examined in a JEOL 1200EX or a TecnaiG^2^ Spirit BioTWIN transmission electron microscope. Images were recorded with an AMT 2k CCD camera.

### Proteomic analysis of EVs by LC-MS/MS

*S. aureus* EVs (8-10 μg) were subjected to SDS-PAGE and stained with Coomassie Blue R-250. Gel sections were analyzed by the Taplin mass spectrometry facility at Harvard Medical School. Peptide sequences (and hence protein identity) were determined by matching protein databases with the acquired fragmentation pattern using the software program Sequest (Thermo Fisher Scientific, Waltham, MA). Proteins were identified by a minimum of two peptides and at least one unique peptide. Sequence analysis was performed with a database containing protein sequences of the *S. aureus* USA300 FPR3757 genome downloaded from NCBIprot. The subcellular localization of each identified protein was predicted by PsortB v.3.0 (www.psort.org/psorb/).

### Real time RT-PCR assay

*S. aureus* strains were cultivated in 5 ml TSB at 37°C to an OD_650 nm_ of 0.9. After centrifugation at 4°C, the bacterial cells were mixed with glass beads in 300 μl lysis buffer (RNeasy mini kit; Qiagen) and lysed by using a high speed Ultramat 2 Amalgamator (SDI, Inc.). Total RNA from the lysate supernatant was purified with the RNeasy mini kit (Qiagen), treated with DNase I (Invitrogen), and stored at −70°C. cDNA was synthesized from 1 μg of bacterial RNA using a Protoscript II First Strand cDNA synthesis kit (New England Biolabs). 50 ng of synthesized cDNA was subjected to Real-time RT-PCR using a Power Green PCR Master Mix (Applied Biosystems) with primers listed in Table S4 and detected in a StepOnePlus RealTime PCR System (Applied Biosystems). The relative transcriptional levels of *hla*_H35L_ and *lukE* were calculated using the ΔΔCt method by normalizing to the 16s rRNA transcriptional level.

### Immunoblotting assays

For Western blots, 10 μg *S. aureus* EVs were subjected to SDS-PAGE, transferred to nitrocellulose membranes, and blocked with PBS + 0.05% Tween-20 (PBST) and 1% skim milk for 1 h at room temperature (RT). After washing with PBST, the membranes were incubated with rabbit anti-LukS-PV (IBT Bioservices) or mouse anti-Hla monoclonal antibody (mAb) 6C12 (IBT Bioservices) overnight at 4°C. The membranes were washed and incubated with secondary antibodies (HRP-conjugated goat anti-rabbit IgG or HRP-conjugated goat-anti mouse IgG) for 2 h at RT before developing the blots using TMB membrane peroxidase substrate (Kirkegaard & Perry Laboratories, Inc). Purified Hla (List Biological Labs) and LukE (IBT Bioservices) were used as positive controls.

For EV dot blotting assays, intact or sonicated EVs were applied to nitrocellulose membranes using a 96 well dot blotter system (Bio-Rad). To block the staphylococcal IgG binding proteins Spa and Sbi, the membranes were blocked with PBST + 5% skim milk and incubated overnight at 4°C with an irrelevant human IgG1 monoclonal antibody (10 μg/ml) in PBST + 1% skim milk. The membrane was washed with PBST and incubated overnight at 4°C with sera (diluted 1:1000 in PBST + 1% skim milk) pooled from mice immunized with EVs (see below) or murine mAbs^78^ to CP5 (4C2; 1.2 μg/ml) or CP8 (5A6; 1.2 μg/ml). After washes with PBST, the membrane was incubated with alkaline phosphatase (AP)-conjugated goat anti-mouse antibody (1:15000 dilution in PBST + 1% skim milk) at RT for 2 h. The membrane was washed with PBST and developed with AP membrane substrate (KPL).

### EV cytotoxicity

The relative toxicity of *S. aureus* EVs (1 to 20 μg/ml) toward human A549 lung epithelial cells, neutrophil-like HL60 cells, and rabbit erythrocytes was assessed. A549 lung epithelial cells grown in a 96-well plate were incubated overnight at 37°C with EVs or 1 μg/ml of purified Hla. Toxicity was assessed using an LDH cytotoxicity assay kit (ThermoFisher Scientific). Differentiated HL60 cells (2 × 10^5^ cells) were seeded in 96-well plate and treated with EVs or 1 μg/ml of Panton-Valentine leukocidin (PVL) for 4 h at 37°C. Cell viability was measured with a CellTiter kit (Promega). A 2% rabbit erythrocyte suspension was mixed with EVs or 1 μg/ml Hla in a 96-well plate for 1 h at 37°C. The erythrocytes were pelleted by centrifugation, and hemolysis was recorded by measuring the OD_545 nm_ of the supernatant using an ELISA reader.

### Animal studies

Mouse experiments were carried out in accordance with the recommendations in the PHS Policy on Humane Care and Use of Laboratory Animals, and animal use protocols were approved by the Partners Healthcare Institutional Animal Care and Use Committee. Female Swiss Webster mice (4 weeks old; Charles River) were immunized by the subcutaneous (s.c.) route on days 0, 14, and 28 with 5 μg/ml of Δ*agr* EVs, Δ*agr*Δ*spa* EVs, or eng-EVs. Control animals were immunized similarly with bovine serum albumin (BSA; Sigma). Blood was collected from the mice by tail vein puncture before each vaccination and again before challenge. Sera were diluted 1.100 and tested by ELISA on 96-well plates coated with 5 μg/ml sonicated WT EVs, 5 μg/ml LukE, or 1 μg/ml Hla. Immunized mice were inoculated with ~2 × 10^8^ CFU *S. aureus* by intravenous (IV) tail vein injection two weeks after the third vaccination. Survival was monitored up to 14 days post-challenge, and the data were analyzed using the log-rank test.

### Toxin neutralization assays (TNAs)

For Hla TNAs, the assay was performed as we previously described^88^. For leukocidin TNAs, human blood was collected from healthy volunteers giving written informed consent, as approved by the Institutional Review Board of The Brigham and Women's Hospital (Human Subject Assurance Number 00000484). Neutrophils were isolated from 10 ml human blood using Polymorphprep (Accurate Chemical), washed, and suspended in RPMI (Invitrogen) containing 5% fetal bovine serum (Invitrogen). Sera from immunized mice were serially diluted and mixed with toxin concentrations yielding ~75% cell lysis (12.5 μg/ml LukED, 2.5 μg/ml PVL, 1 μg/ml HlgAB, or 2 μg/ml HlgCB (1:1 S and F subunits). Samples were pre-incubated with leukocidins for 30 min at RT before the addition of neutrophils (1.2 × 10^5^ cells). After 2h at 37°C in 5% CO_2_, the cells were harvested by centrifugation and suspended in fresh medium. Cell viability was evaluated using CellTiter kit (Promega) according to the manufacturer’s recommendations. Percent neutralization was calculated using the formula: [% Viability of (serum + leukocidin + neutrophils) - % Viability of (leukocidin + neutrophils)].

## Author contributions

X.W. initated the project, and X.W., C.W. and J.C.L designed experiments. X.W. performed experiments. X.W. and J.C.L analyzed data, and X.W., C.W., and J.C.L wrote the manuscript.

## Acknowledgements

We are grateful to Drs. Michael Otto for providing the *S. aureus psm* mutants, Jianxun Ding for providing assistance with DLS and NTA experiments, and Matthew Waldor for use of the StepOnePlus Real-Time PCR System. Christopher Thompson and Anthony Yeh provided expert technical assistance.

## Data availability

Mass spectrometry proteomics data were deposited in the ProteomeXchange Consortium (http://proteomecentral.proteomexchange.org) via the PRIDE partner repository^89^ with the data set identifier PXD007953. Additional data that support the findings of this study are available from the corresponding author upon request.

---

**Figure S1.**
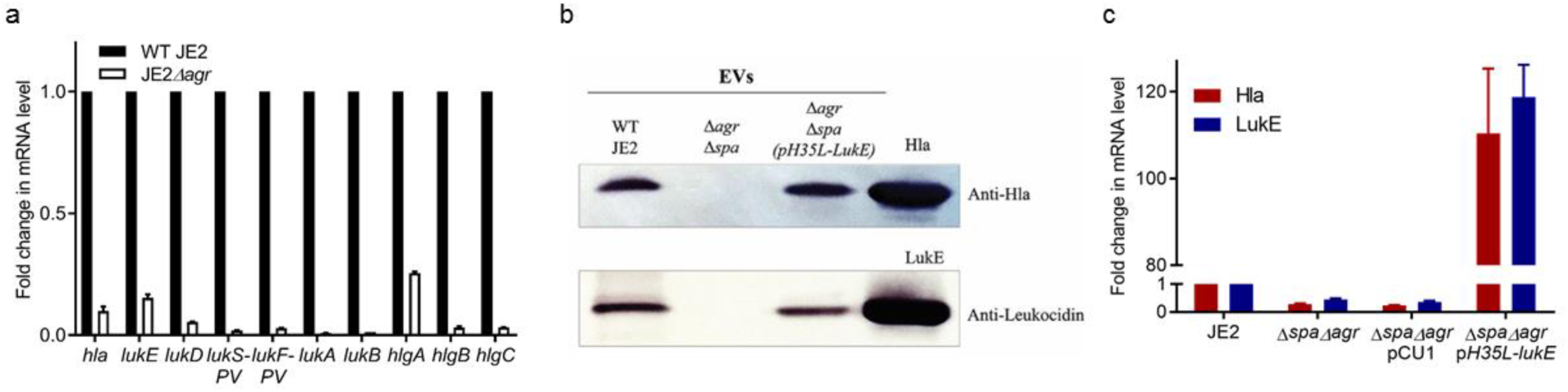
EVs from strain JE2Δ*agr*Δ*spa* (pHla_H35L_-LukE) package recombinant Hla_H35L_ and LukE. **a)** Real time RT-PCR analysis revealed that the mRNA levels of *hla* and genes encoding the leukocidin subunits were dramatically reduced in an *agr* mutant compared to the WT strain JE2. Data are expressed as mean ± SEM relative to the WT strain, and each strain was tested in three replicates. **b)** Purified EVs were subjected to SDS-PAGE. Western blot analysis revealed that Hla_H35L_ and LukE were detected in EVs from recombinant strain JE2Δ*agr*Δ*spa* (pHla_H35L_-LukE) but not in EVs prepared from JE2Δ*agr*Δ*spa*. **c)** Real time RT-PCR analysis revealed that the expression of *hlaH35L* and *lukE* was enhanced ~100-fold in JE2Δ*agr*Δ*spa* (pH35L-LukE) compared to the parental strain JE2. Data are expressed as mean ± SEM, and each group was tested in three replicates.

**Figure S2.**
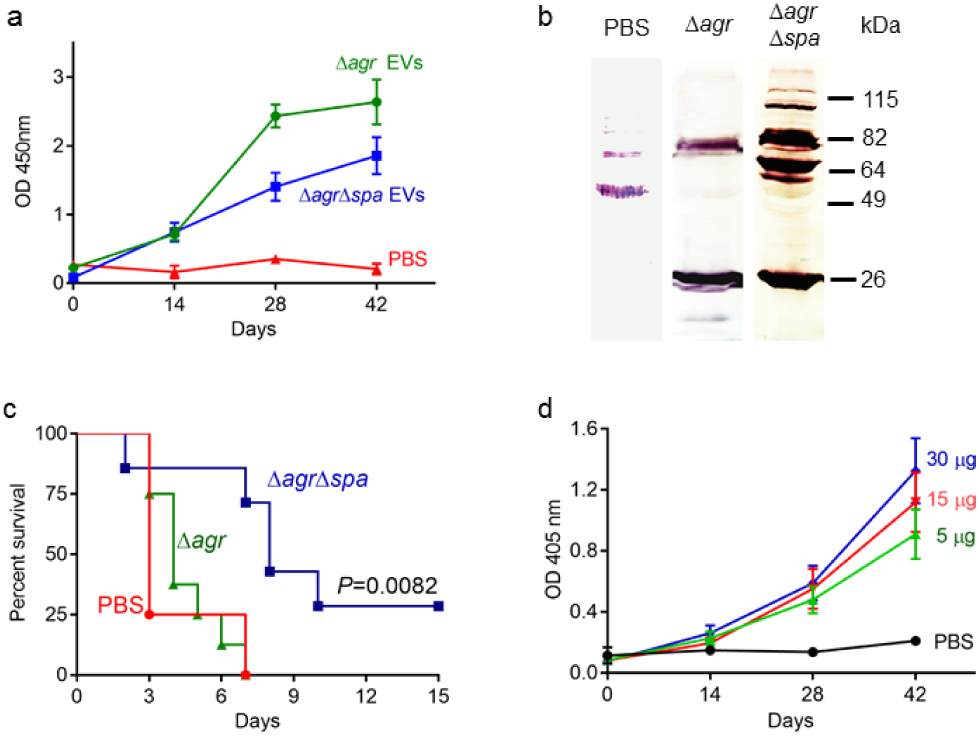
Evaluation of the protective efficacy of *S. aureus* EVs. **a)** Antibody levels in sera (diluted 1:100) from mice immunized with different EVs were analyzed on ELISA plates coated with sonicated JE2 EVs. **b)** USA300 strain FPR3757 cell lysates were subjected to SDS-PAGE. Western blot analysis was performed with sera from mice immunized with different EV preparations. **c)** EV-immunized mice (n=8) were challenged IV with 2×10^8^ CFU strain FPR3757. Mice immunized with EVs were compared to mice given PBS, and survival was analyzed with the log rank test. **d)** Antibody levels in sera (diluted 1:100) from mice immunized with different doses of EVs with alum were analyzed on ELISA plates coated with sonicated JE2 EVs.

**Figure S3.**
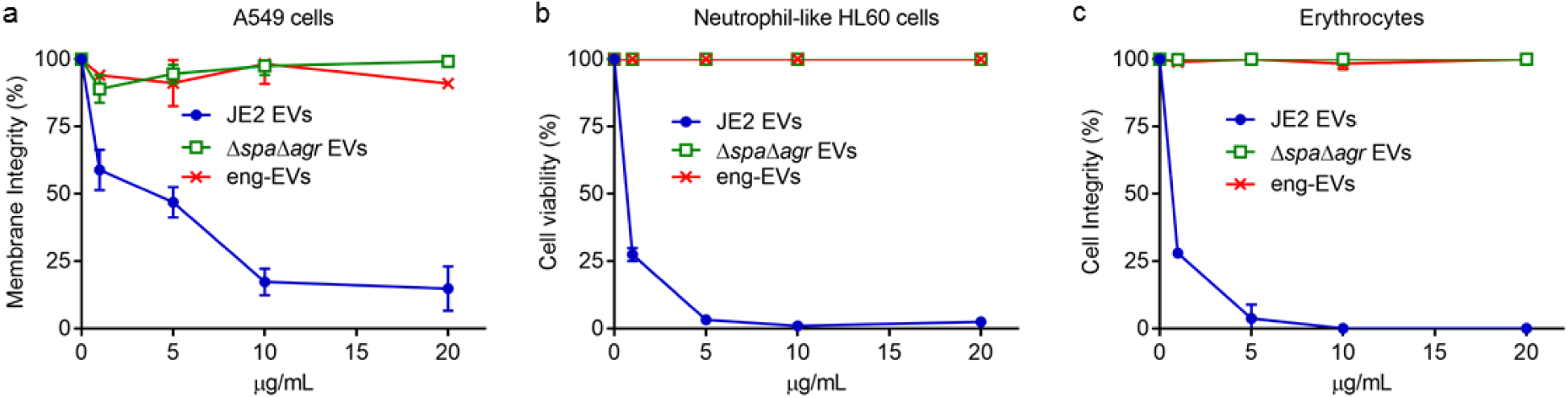
Cytotoxicity of *S. aureus* EVs. **a)** Human lung A549 lung epithelial cells, **b)** neutrophil-like HL60 cells, and **c)** rabbit erythrocytes were treated with different concentration of EVs produced by WT JE2, JE2Δ*agr*Δ*spa*, and eng-EVs.

